# Glycyrrhizic acid improves cognitive levels of aging mice by regulating T/B cell proliferation

**DOI:** 10.1101/2020.03.25.008821

**Authors:** Ruichan Jiang, Jiaming Gao, Junyan Shen, Xiaoqi Zhu, Hao Wang, Shengyu Feng, Ce Huang, Haitao Shen, Hailiang Liu

## Abstract

Licorice is the root of *Glycyrrhiza glabra* L. (Leguminosae), which grows in various warm climates such as the Middle East, Asia, and Southern Europe. It is one of the oldest known medicinal herbs and is referred to as “the father of herbal medicine.” Glycyrrhizic acid (GA) (Fig. 1A), a triterpenoid saponin, is a major component of licorice. It has a variety of pharmacological activities such as anti-inflammatory, antioxidant, anticancer, neuroprotective, and immune effects, among others.^[1]^. Previous studies indicated that GA produces robust neuroprotection via the modulation of anti-apoptotic and pro-apoptotic factors, primarily through the ERK signaling pathway and its anti-inflammatory properties against high-mobility group box 1 phosphorylation and the suppression of inflammatory cytokine induction ^[2-4]^. These results were based on pathological models. Although numerous pathways have been implicated in the neuroprotective effects of GA, the molecular mechanisms are not yet completely understood. In this study, we aimed to investigate the effects of GA in preventing age-related immune involution and cognitive disorders, the relationship between immune involution and cognition, and the underlying molecular mechanisms.

---

Twelve-month-old female C57BL/6 mice were injected with GA (5mg/kg, once every other day, tail intravenous injection, i.v.) for 6 weeks. Then, we tested the behavioral effects of GA using the Morris water maze (MWM) task to evaluate spatial learning and memory. The MWM showed that the GA-treated mouse latencies to the platform gradually decreased compared with the control after six days of training (Fig. 1B). The impairment in learning and memory displayed in middle-aged mice can be prevented by GA treatment as indicated by decreased escape latencies (Fig. 1C), more time in the target quadrant (Fig. 1D), and increased platform crossings (Fig. 1E). This effect was accompanied by improved cerebral blood flow (CBF) (Fig. 1F, G). These data suggest that GA improved CBF and prevented impairments of learning and memory displayed in middle-aged C57BL/6 mice.

We analyzed the transcriptome of C57BL/6 mice blood using RNA-sequencing (RNA-seq). The KEGG pathway showed that GA could significantly influence hematopoietic cell lineage (Fig. 1H). The differential genes in the pathway were clustered using heat mapping (Fig. S1). The RNA-seq results showed that GA activated CD8a, Fcer2a (CD23), and Cr2 (CD21) expression (Fig. 1I) and that it inhibits the expression of macrophage and neutrophil-related genes (Fig. S1). To further verify the relationships of the active genes to cognitive ability, we analyzed the levels of CD3e^+^ positive T cells, CD45R/B220^+^ positive B cells, and CD49b^+^ positive NK cells in the blood and spleen after 22 days of GA treatment. Flow cytometry results showed that GA could increase T and B cell numbers in the blood and spleen (Fig. 1J and K), whereas NK cell numbers were unchanged (Fig. S2).

**Figure 1.**
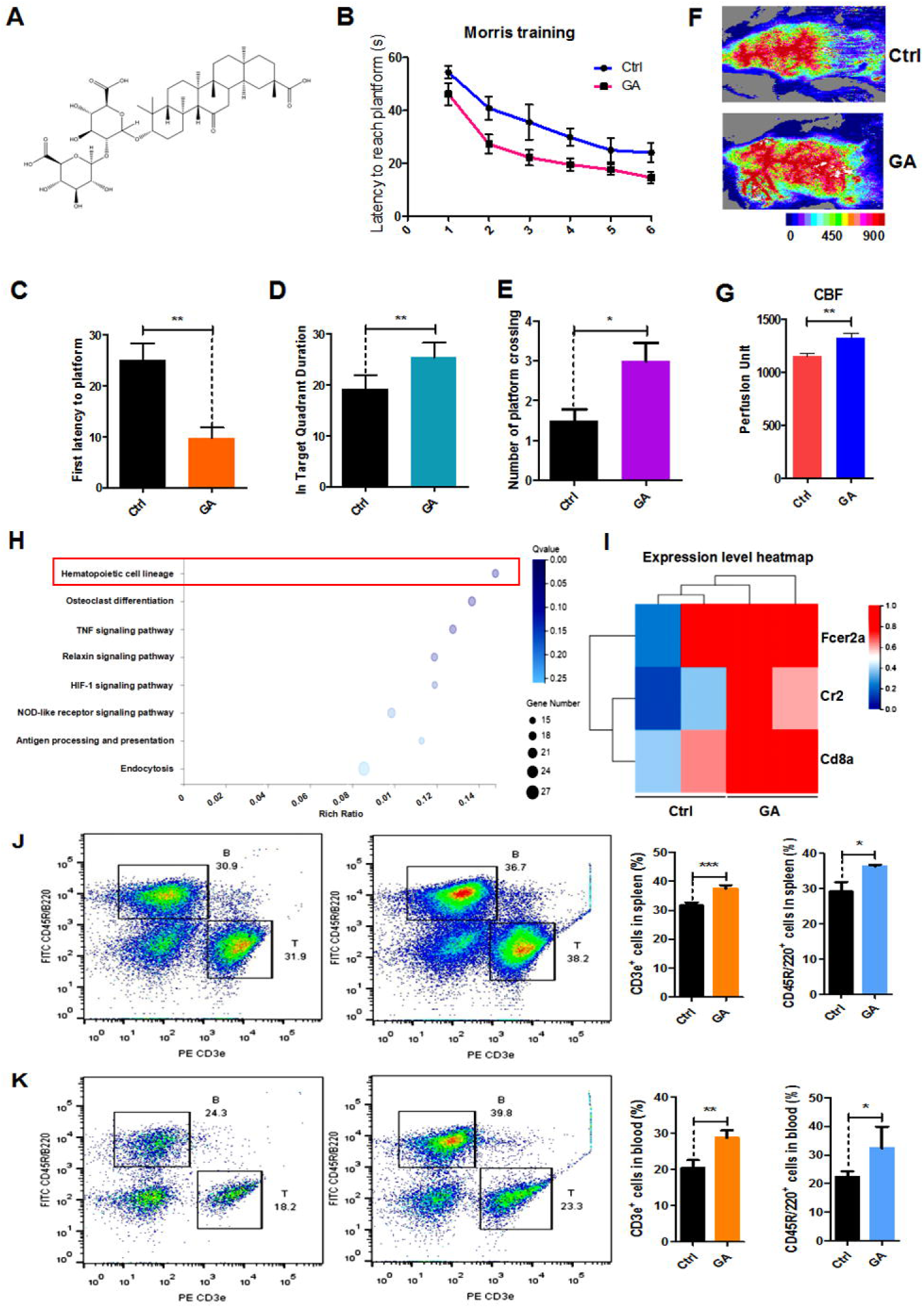
Glycyrrhizic acid ameliorated cognitive declines in aged mice, improved CBF and increased T and B cell numbers. A. The chemical structure of glycyrrhizic acid (GA). B. Escape latencies from six consecutive daily tests. C. The first platform-finding latencies in the probe trial. D. The time spent in the target quadrant in the probe trial. E. The target platform crossing times in the probe trial. F. Images of CBF in C57BL/6 mice. G. Quantification of blood perfusion in the ipsilateral hemisphere. H. Bubble diagram showing GA treatment influence on hematopoietic cell lineage. I. Heatmap showing actively expressed genes related to T cells in the hematopoietic cell lineage of control and GA-treated C57BL/6 mice. J. Representative FACS plots showing CD3e+ T and CD45R/B220+ B cells in the spleens of C57BL mice; bar graphs showing the statistical analysis of T and B cells in the spleens of C57BL mice. K. Representative FACS plots showing CD3e+ T and CD45R/B220+ B cells in the blood of C57BL mice; Bar graphs showing the statistical results of T and B cell analyses in the blood of C57BL mice. GA (5mg/kg, once every other day); Data represent mean ± SEM (n = 8 per group). *P < 0.05, **P < 0.01, ***P < 0.001 vs. Ctrl.

To demonstrate that GA produces the similar effects *in vitro*, we used GA-treated single cells obtained from 8-week-old C57BL mouse spleens for 36h and measured the changes of T and B cells. Flow cytometry showed that GA could significantly increase the numbers of T and B cells *in vitro* (Fig. S3A). Quantitative real-time PCR showed that *CD3e* and *CD45R/B220* genes are overexpressed (Fig. S3B, C). We treated T cells and B cells obtained from 8-week-old C57BL mouse spleens using flow cytometry sorting with 50 µM GA and respectively measured effector T cells (CD45^+^CD3^−^CD45R/B220^+^CD138^+^CD27^+^) and effector B cells (CD45^+^CD3^+^CD44^+^CD62l^−^). The results showed that GA could increase effector cells *in vitro* (Fig. S3D, E). These results show that GA can increase T and B cells *in vivo* and *in vitro*.

To further investigate whether the neuroprotective effects of GA were related to effects on T and B cells, we tested behavioral effects after immune reconstitution (IR) and GA treatment by performing a novel object recognition task to measure the attention and nonspatial declarative memory in B-NDG (B:Biocytogen; N:NOD background; D:DNAPK (Prkdc) null; G:IL2rgknockout) mice. Before the novel object recognition task, we performed the open field test to evaluate the anxiety behavior of these mice. Anxiety levels can be determined based upon the time the mice remain in the corner of the enclosure. We found that the ratio of the time the IR+PBS and IR+GA mice spent in the center of the enclosure to the time spent in a corner over a 10 min test session was not significantly different compared with the control group (Fig. 2A). The novel object recognition task showed that IR+PBS mice exhibited a significant preference for object exploration (Fig. 2B). After IR, the GA treatment showed a more significant preference when contrasted with PBS treatment (Fig. 2B).

**Figure 2.**
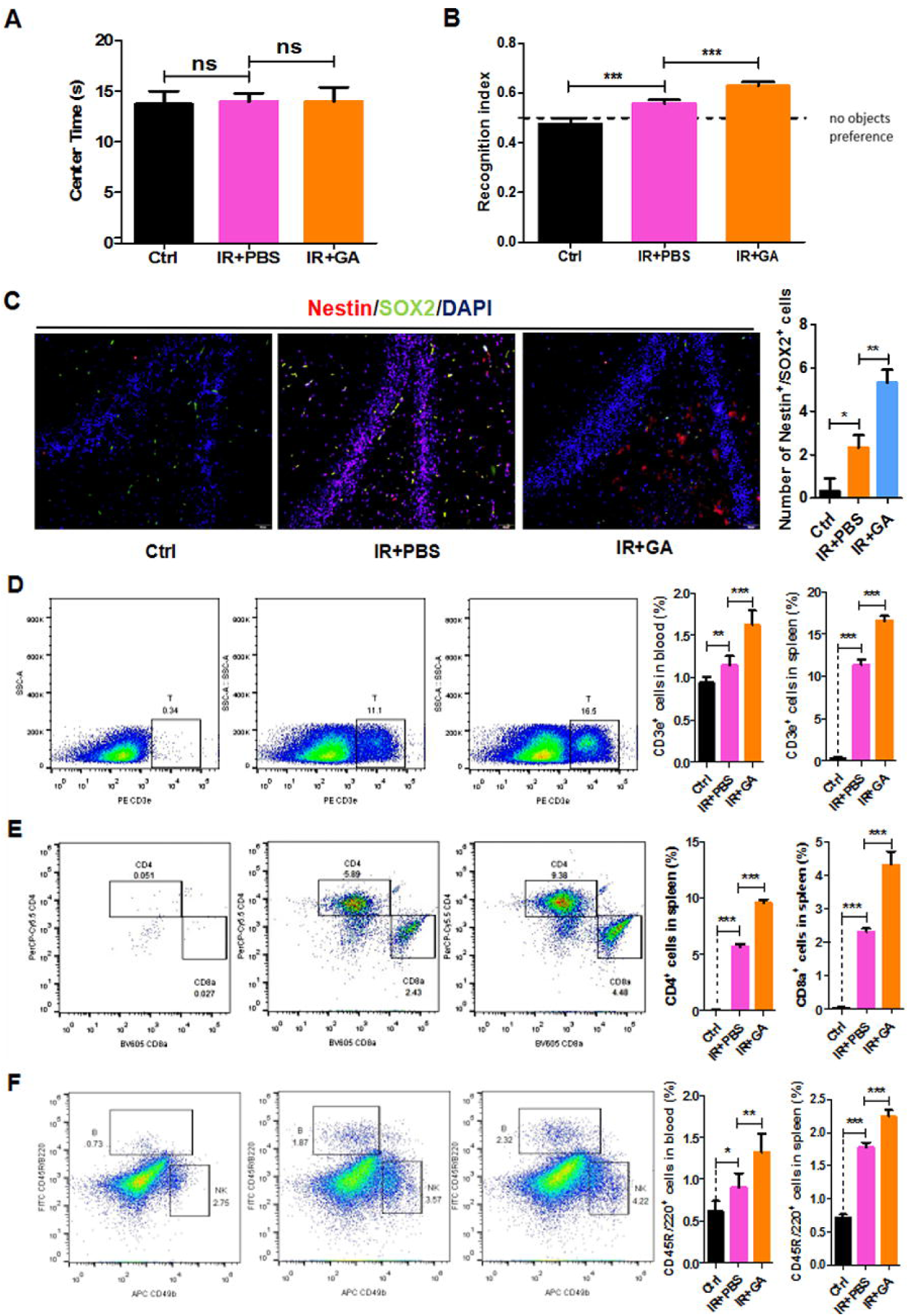
GA treatment increases the speed of IR in B-NDG mice and enhances cognitive ability. A. Anxiety was measured using the open-field test. Results are expressed as the ratio of the percentage of time spent in the center area and the whole experiment time. B. IR+PBS and IR+GA mice showed a preference for novel objects after a 1 h delay between identical object exploration during training and the introduction of a novel object during the test phase. C. Nestin and SOX2 neural stem cell marker expression in B-NDG mouse dentate gyrus. Quantitative analyses of the percentages of nestin+ SOX2+ double-positive cells in the dentate gyrus. D. Representative FACS plots showing CD3e+ T cells in the spleens of B-NDG mice; bar graphs illustrating the statistical results of T cell analyses in the blood and spleens of B-NDG mice. E. Representative FACS plots showing CD4 and CD8 T cells in the spleens of B-NDG mice; bar graphs illustrating the statistical results of CD4 and CD8 T cell analyses in the spleens of B-NDG mice. F. Representative FACS plots showing CD45R/B220+ B and CD49b+ NK cells in the spleens of B-NDG mice; Bar graphs for the statistical results of B cells in the blood and spleen of B-NDG mice. GA (5mg/kg, once every other day); The data represent the mean ± SEM (n = 8 per group). *P < 0.05, **P < 0.01, ***P < 0.001 versus controls.

We analyzed the levels of CD3e^+^ positive T cells, CD45R/B220^+^ positive B cells, and CD49b^+^ positive NK cells in the blood and spleen. Flow cytometry results showed that the IRs of B-NDG mice were successful given that T, B, and NK cells in IR+PBS mice were significantly increased compared with controls (Fig. 2D, F, Fig. S4). Thus, GA treatment after IR can increase the number of T and B cells compared with PBS treatment after IR (Fig. 2D, F), whereas NK cell numbers were unchanged (Fig. S4). This result is consistent with those above from C57BL mice. Furthermore, we measured the numbers of CD4 and CD8 cells in the spleen and found they were increased (Fig. 2E), which was consistent with our RNA-sequencing results.

To further investigate various mouse brain tissue characteristics, we examined the neural stem cell markers nestin and SOX2 in the dentate gyrus (DG) of the hippocampus in IR+PBS, or IR+GA treated mice and control mice. The results showed that the numbers of nestin+ Sox2+ double-positive cells were significantly increased in the DG of immunologically reconstituted mice compared with the control and GA treatment, or compared with PBS injection after IR (Fig. 2C).

Our work has thus revealed that GA treatment can revitalize the aged brain and alleviate aging-associated cognitive declines and that these effects may be related to the immune system. RNA-seq data demonstrated the effect of GA on hematopoietic cell lineage, and we posited that GA could enhance cognition through immune system modulation. We observed lymphocyte changes in C57BL mice, then used B-NDG mice to verify further the importance of the immune system in cognitive ability. The B-NDG mice exhibit a severe immunodeficiency phenotype with no mature T cells, B cells, or functional NK cells and a lack of cytokine signaling ability. The results of the novel object recognition task confirmed our hypothesis; after IR, cognitive ability was improved (Fig. 2B). Although we cannot confirm which groups of cells are essential in the associated mechanisms, these results reveal a secure connection between the immune system and cognition.

A marked increase in the number of cytotoxic CD4 T cells (CD4 cytotoxic T lymphocytes) is a signature characteristic of supercentenarians ^[5]^. We observed that CD4^+^ T cells are significantly expanded in number (Fig. 2E), but we cannot confirm whether these represent cytotoxic CD4 T cells. The potential effects of GA treatments on memory and behavior, and even longevity, may be worthy of future investigation.

The brain is particularly susceptible to the effects of aging, and aging and associated inflammation is a major risk factor for a variety of neurocognitive and neurodegenerative diseases ^[6]^. There are abnormal neural stem cells, neurons, and microbes in the brain, and their clearance by immune cells can preserve cognitive function ^[7]^. Further efforts to understand the effector cells in this process and how these events occur would be worthwhile in understanding the roles of immune cells in brain aging.

During cytokine storms, cytokine levels are abnormally high, which can lead to fever, low blood pressure, and heart problems and, in some cases, organ failure and death ^[8]^. Bacterial infection and viruses such as SARS and MERS can cause cytokine storms, which attack host cells, mainly the patient’s immune cells ^[9]^. Macrophages and neutrophils are known to produce catecholamines in response to inflammatory stimuli such as lipopolysaccharide (LPS), which is a hallmark of many types of bacterial infection. ^[10]^. Our RNA-seq results showed that GA treatment could inhibit several genes related to macrophages, neutrophils, and IL1β (Fig. S1). This finding indicates that GA might inhibit the proliferation of macrophages and neutrophils from preventing cytokine storms. Treatment with GA can also alleviate β-amyloid ^[^11, 12^]^ or systemic LPS-induced ^[13]^ cognitive impairment via inhibition of neuroinflammation. Some COVID-19 patients have been cured using diammonium glycyrrhizinate, an ammonium salt preparation of 18-alpha-GA. This phenomenon could be explained by the anti-inflammatory properties of GA. In conclusion, the potential effects of GA treatments on memory and behavior, longevity, and anti-viral activity may be worthy of future investigation.

## Acknowledgements

This work was supported by grants from the National Key Research and Development Program (2019YFC200187), National Natural Science Foundation of China (Grant No 31671539, 31370214), and Major Program of Development Fund for Shanghai Zhangjiang National Innovtaion Demonstration Zone<Stem Cell Strategic Biobank and Stem Cell Clinical Technology Transformation Platform> (ZJ2018-ZD-004), and the Oasis Scholars Support Program of Shihezi University (for Liu HL).

