## Supplementary Materials for "Glycyrrhizic acid improves cognitive levels of aging mice by regulating T/B cell proliferation"

**Materials and methods**

**Reagents**

Glycyrrhizic acid (GA) was obtained from TAUTO (Sichuan, China). 4',6-diamidino-2-phenylindole dihydrochloride (DAPI), lysis buffer, and all antibodies used for flow cytometry were purchased from BD Pharmingen (San Diego, CA). Information regarding the antibodies used in this study is listed in the Resources Table.

**Animals**

Both 8-week-old and 8-month-old C57BL/6 mice were obtained from Shanghai SLACCAS Co., Ltd (Shanghai, China). Eight-week-old NOD-SCID IL-2 receptor gamma null mice were obtained from Jiangsu Biocytogen Co., Ltd. (Nantong, China). All mice were housed five per cage and maintained on a 12 h light/dark schedule and allowed free access to food and water following a protocol approved by the Animal Research Committee of Tongji University School of Medicine, China.

**qRT-PCR analysis**

Total RNA was isolated using TRIzol reagent (Thermo Fisher Scientific,Waltham, MA, USA), and cDNA was prepared using the Prime Script™ RT Master Mix (Perfect Real Time) (Takara, Dalian, China) according to the manufacturer's protocols. The qRT-PCR reactions were performed using SYBR green fluorescent dye (BioRad). Primer sequences are listed in the Resources Table.

**The Morris water maze task**

The Morris water maze consists of a round, black pool of 120 cm in diameter and 31 cm deep containing water at 23 ± 1 °C. The escape platform (11 cm in diameter, adjustable height) was placed in the center of one quadrant of the pool and hidden below the water surface at 1.5 cm deep. Various prominent visual cues were placed around the pool and remained in the same position during training and testing periods. Each group was trained for 6 consecutive days from four locations and then tested on day 6 with a new direction, removing the hidden platform and allowing free swimming for 60 s. The escape latencies from the water, and the distance traveled to find the platform was recorded using video-animal tracking software.

**Novel object recognition**

Each mouse was habituated to an empty novel object recognition (NOR) open-field box for two 10-min test sessions 24 h apart. Twenty-four hours after the last habituation session, mice were subjected to training during a 10 min exposure session of two identical, nontoxic objects (metal or hard plastic items) in the open-field box. The time spent exploring each object was recorded using ObjectScan software (Clever Sys. Inc, Reston, VA); an area 2 cm^2^ surrounding the object was defined, so that nose entries within 2 cm of the object was recorded as time exploring the object. After the training session, the animal was returned to its home cage. After a retention interval of 1 h, the animal was returned to the arena in which two objects, one identical to the familiar object but previously unused (to prevent olfactory cues and prevent the necessity of washing objects during experimentation) and one novel object. The animal was allowed to explore for 10 min, during which the amount of time exploring each object was recorded. Objects were randomized and counterbalanced across animals. Animals that spent <7s exploring the objects during the 10 min test session were omitted from the analysis. Objects and arenas were thoroughly cleaned with 70% isopropanol between trials.

**Open field test**

The device is based on a square field (50 × 50 × 30 cm). A lamp was located 150 cm above the field, and the illumination of the central area was approximately 100 lux. At the beginning of each experiment, a mouse was placed in a 15 × 15 cm central area. At the beginning of the experiment, each mouse was placed in the center of the field, and the time spent in the central area within 10 min was recorded.

**Cortical cerebral blood flow measurements**

Images were acquired with a laser speckle contrast imager (PeriCam PSI System, Perimed, Stockholm, Sweden). We used the PeriCam PSI HD system to calculate an arbitrary index of cerebral blood flow (perfusion units) in the ipsilateral hemisphere.

**Cell preparation and flow cytometric analysis**

Harvested mouse spleens were macerated and washed with phosphate buffer saline (Thermo Fisher Scientific,Waltham, MA, USA). Then, splenocytes were obtained by removing the red blood cells with lysis buffer (BD Biosciences, San Jose, CA, USA) and filtering through a 70 μm cell strainer (Jet Biofil, Guangzhou, China). Additionally, fresh blood samples were collected in heparinized tubes. T (CD3+) cells, B (CD3-CD45R+) cells, and NK (CD3-CD49b+) cells in the spleen and blood were directly quantified using flow cytometry (Beckman FC-500, Miami, FL, USA).

Some splenocytes were cultured with added GA in 6-well plates for 36 h. Then effector T (CD45+CD3+CD44+CD62l-) cells, and effector B (CD45+CD3-CD45R+CD138+ CD27+) cells were directly quantified using flow cytometry.

**Immunofluorescence images**

Brain tissue was collected from mice following treatment with GA. Brain tissue was fixed with 4% paraformaldehyde (PFA) in 0.1M phosphate-buffered saline (PBS) overnight, followed by a 15-30% sucrose gradient dehydration for one day until the brain had completely sunk to the bottom of the sucrose solution. An optimal cutting temperature compound was used to embed tissue samples. Successive coronal sections of 10 μm were cut using a freezing microtome. Tissue slices were washed with PBS and blocked for 1h (in 10% bovine serum albumin, 3% normal donkey serum, and 1% Triton X-100 in PBS) and mounted on tissue slides. Antibodies against Nestin, SOX2 (Abcam, Cambridge, MA, USA). The next day, the slides were washed three times and incubated with the appropriate Alexa 488- and Alexa 568- secondary antibodies (Thermo Fisher Scientific) for 1 h at room temperature (1:1,000 dilution). DAPI staining was used to label nuclei. Slides were examined using an OLYMPUS BX53 microscope (Olympus, Madison, WI).

**RNA sequencing analysis**

RNA sequencing was performed independently and uniformly for each sample. GA- and control-treated mice were anesthetized and euthanized, and two blood samples were removed for RNA-seq following extraction of total RNA. Clean reads were aligned to the reference gene sequence using bowtie-2, and the gene expression levels of each sample were calculated. DEG detection was conducted using the DEGseq method. The statistical results were based on the ma-plot method. The number of reads of specific genes obtained from the sample was sampled randomly, and then *P*-values were calculated according to the normal distribution and corrected to *q*-values. To improve the accuracy of DEG detection, genes with a difference multiple of more than twice, and a *q*-value of ≤0.001 were screened and defined as significantly differentially expressed genes. The RNA-seq data files generated in this study are available in the NCBI Gene Expression Omnibus (GEO) under the accession number GEO: GSE146239.

**Statistical analysis**

Statistical analysis of data was conducted using Graphpad Prism 5.0 and expressed as the mean ± standard error of the mean (SEM). Statistical comparisons of two groups were made using the unpaired *t*-test. Probability values less than 0.05 were considered statistically significant.

**Resources table**

| **Reagent or Resource** | **Source** | **Identifier** |
| --- | --- | --- |
| **Antibodies** |  |  |
| CD3e | BD Biosciences | Cat# 553063 |
| CD45R/B220 | BD Biosciences | Cat# 553087 |
| CD49b | BD Biosciences | Cat# 558295 |
| CD4 | BD Biosciences | Cat# 550954 |
| CD8a | BD Biosciences | Cat# 563152 |
| CD45 | BD Biosciences | Cat# 566439 |
| CD3 | BD Biosciences | Cat# 560590 |
| CD44 | BD Biosciences | Cat# 553134 |
| CD62l | BD Biosciences | Cat# 560516 |
| CD45R/B220 | BD Biosciences | Cat# 553087 |
| CD138 | BD Biosciences | Cat# 558626 |
| CD27 | BD Biosciences | Cat# 558754 |
| **Chemicals** | | |
| Glycyrrhizic acid | TAUTO | Cat# 1405-86-3 |
| Lysing Buffer | BD Biosciences | Cat# 555899 |
| DAPI | BD Biosciences | Cat# 564907 |
| TRIzol | Gbico | Cat# 15596018 |
| the Prime Script™ RT Master Mix (Perfect Real Time) | Takara | Cat# RR036A |
| SYBR Green fluorescent dye | BioRad | Cat#172-5120 |
| PBS | Gbico | Cat# 10010049 |
| PFA | Sigma | Cat# 158127 |
| Nestin | Millipore | Cat# MAB5326 |
| SOX2 | Abcam | Cat#ab93689 |
| **Experimental Model** | | |
| B-NDG | Biocytogen | B-CM-002 |
| C57BL/6 | SLACCAS |  |
| **Oligonucleotides** | | |
| *CD3e*-qPCR-F:  CTGCTACACACCAGCCTCAA | This paper | N/A |
| *CD3e*-qPCR -R:  GTAATAAATGACCATCAGCAAGC | This paper | N/A |
| *CD45R/B220*-qPCR -F:  CCAGTGATGCTACCACAACG | This paper | N/A |
| *CD45R/B220*-qPCR -R:  CAATCCTCATTTCCACACTTAGC | This paper | N/A |
| *β-actin*-qPCR -F:  TATTGGCAACGAGCGGTTC | This paper | N/A |
| *β-actin*-qPCR -R:  ATGCCACAGGATTCCATACCC | This paper | N/A |

**Supplementary Figures**

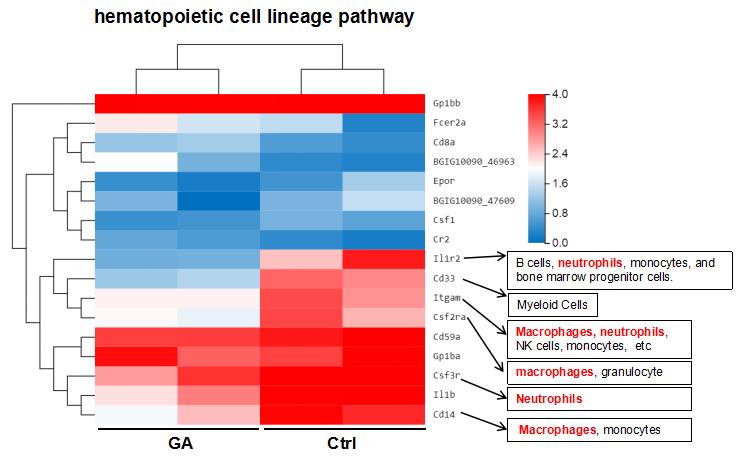

**Figure S1 (Related to Figure 1). Heatmap showing differentially expressed genes (DEGs) in hematopoietic cell lineage of control and GA-treated C57BL/6 mice.**

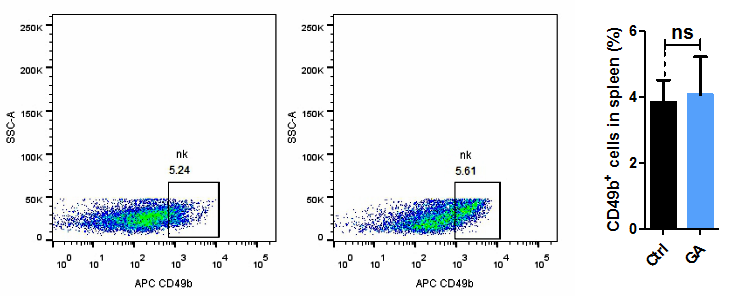

**Figure S2. NK cell numbers were unchanged after GA treatment (Related to Figure 1).** Representative FACS plots showing CD49b+ NK cell numbers in the spleens of C57BL mice; bar graphs show the statistical results for NK cell analyses in the blood and spleens of C57BL mice.

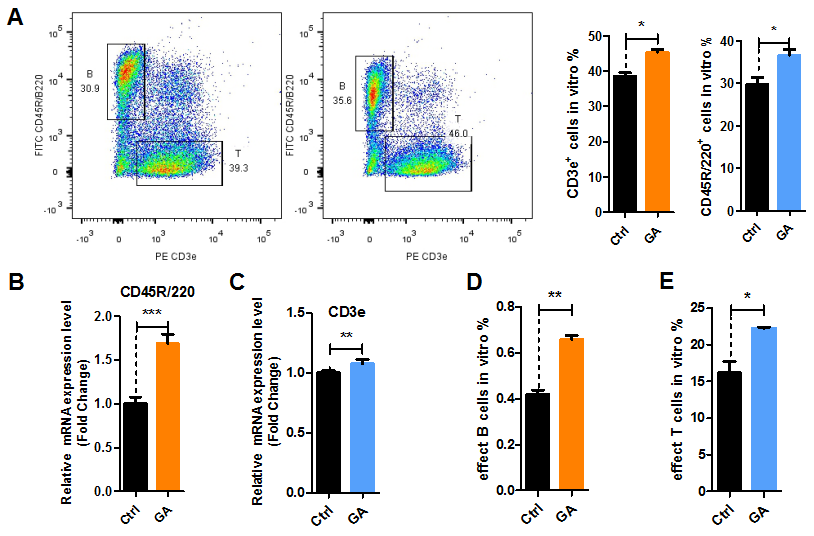

**Figure S3. GA treatment increases the proliferation of T and B cell subsets *in vitro* (Related to Figure 1).** A. Representative FACS plots showing CD3e+ T and CD45R/B220+ B cells in vitro; Bar graphs show the statistical results of T and B cell analysis *in vitro*. B. CD3e mRNA expression in the spleens of C57BL mice. C. CD45R/B220 mRNA expression in the spleen of C57BL mice. D. Bar graphs showing the statistical results of effector B cell analysis after flow cytometry sorting. E. Bar graphs showing the statistical results of effector T cell analysis after flow cytometry sorting. Data represent the mean ± SEM (n = 8 per group). *P < 0.05, **P < 0.01, ***P < 0.001 vs. control.

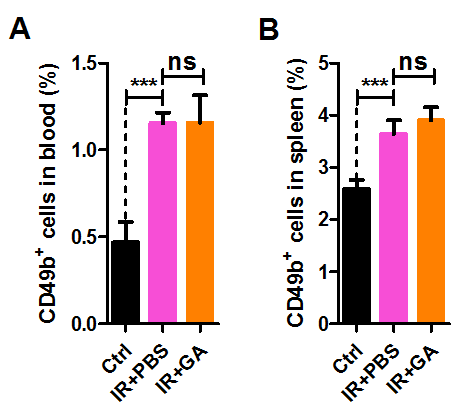

**Figure S4 (Related to Figure 2). Bar graphs illustrating the statistical results of NK cell analyses in the blood and spleen of B-NDG mice.**
